# Multivariate Analysis for Agro-Morphological and Quality Traits in Groundnut (*Arachis hypogaea* L.) Genotypes in Eastern Ethiopia

**DOI:** 10.1101/2025.05.22.655537

**Authors:** Desu Beriso, Seltene Abadi, Abdi Mohammed

**Author notes:** Corresponding author Gmail.

## Abstract

It is imperative to study the level of genetic variability available in the existing groundnut genotypes due to the fact that, the aim of groundnut breeding programs across the world is to develop new varieties that meet the needs of growers, processors, consumers, and overall market demands. The present study was carried out to assess the extent of genetic variability among groundnut genotypes for agro-morphological and quality traits. Thirty-six groundnut genotypes were evaluated in a 6 x 6 simple lattice design during 2023 post-rainy season under irrigation at Dire Dawa, the research station of Haramaya university, Ethiopia. Data were collected on kernel yield and other morphological traits, oil content and oil yield. The data on traits were subjected for principal component (PC) values, clustering and Euclidean distance. In this study, the first six Principal Components Analysis (PCA) found to be significant and accounted for 74.51% of the total variation in which the first principal component (PC1) and the second principal component (PC2) contributed more to the variation. Those 36 genotypes were grouped into six major clusters and the dendrogram showed that cluster I, II, III, IV, V and VI included 6, 9, 8, 5, 7 and 1 numbers of genotypes in that order. Euclidean distance ranged from 2.45 to 8.54 with the mean, standard deviation and coefficient of variation of 5.44, 1.17 and 21.56%, respectively. Based on the result of the current study, there were variations of genetic distances among genotypes, Gv17 and Gv28, Gv3 and Gv23, Gv3 and Gv30, Gv15 and Gv17, Gv22 and Gv28, and Gv3 and Gv34 which could be exploited through hybridization for cultivar development in groundnut breeding programs in Ethiopia. Therefore, these genotypes are recommended as excellent candidates for further breeding and variety development.

## 1. Introduction

Groundnut or peanut (*Arachis hypogaea* L.) is one of the important oilseed crops grown in tropical and subtropical regions of the world (Olayinka and Etejere, 2015). It is an annual legume in the family of *Leguminaceae* native to South America, Mexico and Central America and successfully grown in diverse environments in six continents (Seetha *et al.,* 2018). It ranks 5^th^ among oilseed crops after oil palm, soybean, rapeseed, and sunflower in terms of volume of production globally and widely grown in more than 110 countries of tropical, subtropical, and warm temperate regions (FAOSTAT, 2022). It is a multi-purpose crop largely cultivated in tropical and sub-tropical parts of the world (Olayinka and Etejere, 2015). Africa, Asia and the United States of America produce about 13.78%, 49.84% and 32.35% of the world’s groundnut, respectively (FAOSTAT, 2022).

Globally, China, India, Nigeria, USA and Sudan are the leading groundnut-growing countries (FAOSTAT, 2022). Globally, the estimated annual groundnut production was about 536,389.32 metric ton from 31.6 million hectares of production area. The average global yield was about 1.70 tons per hectare (FAOSTAT, 2022). Similarly, Nigeria, Sudan, Chad, Cameroon, Senegal, Guinea, Burkina Faso and Niger are the leading groundnut producing countries in Africa. In Africa, the annual production of groundnut was 168,602.72 metric ton from 17,430,165 hectare of production area with productivity of 0.97 tons per hectare, while 20,644.29 metric ton from 2.8 million hectare of production area with average yield of 0.74 tons in Eastern Africa (FAOSTAT, 2022).

Groundnut seed is a rich source of oil (35-56%), protein (25-30%), carbohydrates (9.5-19.0%), minerals (Ca, K, Mg and P) and the vitamins (B, E, and K) (Gulluoglu *et al.,* 2016; Amare *et al.,* 2017). The seed oil is useful for determining shelf life, nutrition, stability and flavor of food products (Bertioli *et al.,* 2019). However, seed oils differ in quantity and the fatty acid composition (Bera *et al.,* 2019). The oleic acid is the dominant fatty acid, followed by linoleic acid. The higher oleic acid the better shelf life and the oxidative stability of groundnut products and helps to minimize the risk of cardiovascular disease (CVD), while consumed (Bera *et al.,* 2019). The oleic acid (35.7 to 82.2%) to linoleic acid (2.9 to 40.3%) ratio is varied with seeds and used as a stability and shelf-life index for industrial applications (Bertioli *et al.,* 2019).

Groundnut has a number of industrial purposes manufacturing various products, such as packs of sweets and chocolates as food, as well as used for productions of paints, biofuels, lubricants and insecticides (Variath and Janila, 2017).

In Ethiopia, processed groundnuts are used in diversified ways, such as groundnut butter, which is used as spread for bread or biscuits; in cookies, sandwiches, candies and frostings or icings (Chala *et al.,* 2014) and as groundnut cake, locally called “*Halawa”* packets of groundnut and sauces. It is also used as a substitute for milk in the preparation of *"Makiyato" or “white coffee”* during fasting days (Chala *et al.,* 2014). The groundnut seeds are also directly eaten raw as vegetables and used daily as roasted “ocholonie” or *“Kolo”*; used to prepare children’s food *“fafa”*. As a ‘plumpy’ nut (a peanut based nutritional product), or ready- to- use therapeutic food (RUTF), is also used as a source of nutrition for malnourished children in developing countries including Ethiopia (Getahun *et al.,* 2018). Further, groundnut is an ideal crop in crop rotational systems to improve soil fertility due to its natural capability to fix atmospheric nitrogen (Jaiswal *et al*., 2017). In tropical Africa, groundnut haulms are used as animal fodder and the shells as source of fuel and fertilizer (Ajeigbe *et al.,* 2018), which is also common in eastern lowland parts of Ethiopia

The lowland areas of Ethiopia have considerable potential for increased oil crop production, amongst which groundnut is the second most important lowland oilseed crop after sesame in the country (Amare *et al.,* 2017). The estimated annual groundnut production in Ethiopia was about 205,068.65 tons from 113,514.95 hectares of production area with the productivity of 1.807 tons per hectare (CSA, 2021). In the country, groundnut production is concentrated in some areas of Oromia, which accounted for (59.2%) of the total national production, Benishangul-Gumuz (24.83%), Amhara (7.43%), Harari (3.29%) and SNNP (1.29%) (CSA, 2021). Eastern Hararghe Zone holds primary position in producing and supplying groundnut to domestic markets as compared to other parts of the country (CSA, 2021). Several factors constrained groundnut production in Ethiopia. These include abiotic and biotic stresses, socioeconomic factors (Seltene *et al.,* 2019a). Groundnut production mostly constrained by drought stress, mainly occurring during the flowering, leads to soil-borne fungal infections and poor post-harvest handling and mold development, subsequently aflatoxin productions, which may lead to rot domestic and international market rejects due to maximum aflatoxin accumulations or contaminations. Poor soil fertility, lack of access to improved seed, pre-harvest foliar diseases, use of low yielding varieties, inadequate access to extension services, credit, and availability of resilient varieties. Perhaps, groundnut growers preferred traits including high-shelled yield, early maturity, tolerance to drought stress, high market value, good seed quality, adaptability to local growing conditions, and resistance to root diseases and invasive weed species (Seltene *et al.,* 2019b).

Generation of high yielding resilient varieties with market-preferred or oriented traits is a priority for addressing groundnut yield gap. Genetic variability acts as a basis for development of such improved cultivars upon which selection thrives (Govindaraj *et al.,* 2015). Genotypic and phenotypic variation, genetic advance, path analysis based on genotypic and phenotypic correlations coefficient and heritability have been reported for different traits in groundnut (Rao *et al.,* 2014; Zekeria *et al.,* 2017b, Fantaye *et al.,* 2018). The coefficients of variation provide a basis to compare diversity of quantitative traits while high heritability and genetic advance suggest possibility of effective phenotypic selection (You *et al.,* 2016). These traits indicate the genetic potential of a given germplasm that dictates the success in breeding program (Shrestha, 2016).

Information on the nature and degree of genetic diversity helps to plant breeders in choosing the diverse parents for hybridization (Singh and Nigam, 2016). For a successful breeding program, the presence of genetic diversity and variability play a vital role. Genetic diversity is essential to meet the diversified goals of plant breeding such as breeding for increasing yield, wider adaptation, desirable quality, pest and disease resistance for sustainable groundnut production. Selection of genetically diverse parents in any breeding program is of immense importance for successful recombination breeding (Rivière *et al.,* 2015). Improvement in yield and quality is likely achieved by selecting genotypes with desirable trait combinations existing in the nature or by hybridization. The parent identified on the basis of the divergence analysis would be more promising than other analyses (Singh *et al.,* 2013).

Naturally, groundnut has a narrow genetic base as a result of its monophyletic origin, self-pollination and lack of gene flow, due to origin of the crop through a single hybridization event between two diploid species, followed by a chromosome doubling and crossing barriers with wild diploid species (due to ploidy differences) (Kochert *et al*., 1996). Moreover, to improve and sustain the yield of groundnut, plant breeders should have a better understanding of the genetic variability of yield and its components and development of high yielding cultivars (Bertioli *et al.,* 2019).

Hence there is a need to study the genetic variability of plants for the efficient management and the conservation of races and their optimum utilization in plant breeding. Genetic variability is essential for initiating an effective and successful breeding programs; thus, it is imperative to study the level of genetic variability available in the existing groundnut genotypes grown in Ethiopia. Several studies have been conducted on groundnut in the country particularly in East Hararghe; however, there is limited information on genetic variability, association among morpho-agronomic and quality traits which remains a major concern for groundnut improvement programs. Therefore, the objectives of the current study were to assess the extent and pattern of variation of groundnut genotypes for agro-morphological and quality traits in eastern Ethiopia.

## 2. Materials and Methods

### 2.1. Description of Study Area

The current study was carried out at Dire Dawa, Tony farm Research sub-Station of Haramaya University during 2023 post-rainy season under irrigation. The location is situated at 9° 35’ 0” North and 41° 52’ 0" East at an altitude of 1197 meters above sea level (m.a.s.l) in the eastern Rift Valley escarpment of Ethiopia. The average maximum and minimum temperatures of the area are 31.89 °C and 17.08 °C, respectively and mean annual temperature of 25.36 °C for the period of 1955 to 2015. The area receives average rainfall of about 637 mm per annum. Soils of the farm are predominantly sandy clay loam with pH varies with soil depth which is about 7.97 at a depth of 0 to 20 cm and about 8.44 at a depth of 20 to 40 cm. The bulk density range varies from 1.28 to 1.39 g cm ^-3^. The area is categorized as hot to warm arid according to agro-ecological belts of Ethiopia.

### 2.2. Experimental Materials

Thirty-six groundnut genotypes, including two standard checks were obtained from the National Groundnut Program based at Haramaya University (Table 1).

**Table 1.** List of groundnut genotypes used in this study.

| Codes for Genotype | Genotypes | Breeding status | Seed source |
| --- | --- | --- | --- |
| Gv1 | Babile-1 | Released variety | Haramaya University |
| Gv2 | Babile-2 | Released variety | Haramaya University |
| Gv3 | ICGV 00350 | ABL | ICRISAT-India |
| Gv4 | ICGV 02271 | ABL | ICRISAT-India |
| Gv5 | ICGV 03179 | ABL | ICRISAT-India |
| Gv6 | ICGV 03181 | ABL | ICRISAT-India |
| Gv7 | ICGV 03196 | ABL | ICRISAT-India |
| Gv8 | ICGV 03921 | ABL | ICRISAT-India |
| Gv9 | ICGV 07356 | ABL | ICRISAT-India |
| Gv10 | ICGV 07947 | ABL | ICRISAT-India |
| Gv11 | ICGV 13825 | ABL | ICRISAT-India |
| Gv12 | ICGV 13833 | ABL | ICRISAT-India |
| Gv13 | ICGV 13834 | ABL | ICRISAT-India |
| Gv14 | ICGV 14840 | ABL | ICRISAT-India |
| Gv15 | ICGV 14858 | ABL | ICRISAT-India |
| Gv16 | ICGV 14876 | ABL | ICRISAT-India |
| Gv17 | ICGV 55437 | ABL | ICRISAT-India |
| Gv18 | ICGV 86024 | ABL | ICRISAT-India |
| Gv19 | ICGV 86124 | ABL | ICRISAT-India |
| Gv20 | ICGV 89104 | ABL | ICRISAT-India |
| Gv21 | ICGV 91284 | ABL | ICRISAT-India |
| Gv22 | ICGV 91315 | ABL | ICRISAT-India |
| Gv23 | ICGV 92302 | ABL | ICRISAT-India |
| Gv24 | ICGV 93305 | ABL | ICRISAT-India |
| Gv25 | ICGV 94379 | ABL | ICRISAT-India |
| Gv26 | ICGV 94434 | ABL | ICRISAT-India |
| Gv27 | ICGV 96801 | ABL | ICRISAT-India |
| Gv28 | ICGV 96826 | ABL | ICRISAT-India |
| Gv29 | ICGV 96909 | ABL | ICRISAT-India |
| Gv30 | ICGV 97094 | ABL | ICRISAT-India |
| Gv31 | ICGV 97097 | ABL | ICRISAT-India |
| Gv32 | ICGV 97188 | ABL | ICRISAT-India |
| Gv33 | ICGV 98088 | ABL | ICRISAT-India |
| Gv34 | ICGV 98100 | ABL | ICRISAT-India |
| Gv35 | ICGV 99240 | ABL | ICRISAT-India |
| Gv36 | ICGV 99241 | ABL | ICRISAT-India |

ABL = Advanced breeding line, Gv=Groundnut genotype.

### 2.3. Experimental Design and Field Management

The field experiment was conducted using irrigation during the 2021 post-rainy season. The experiment was laid out in simple lattice design. Seeds of each genotype were sown in 3 rows of 3 meter long with 0.6, 0.1, 0.5 m between rows, plants and plots in that order. Hence, a total of 5.4 m^2^ area was allocated for each experimental plot in each incomplete block within replication. Spacing between sub-blocks (incomplete block) and replications was 0.7 and 1 m, respectively. All agronomic practices were applied as per the standard procedure for groundnut production (Janila *et al*., 2018).

### 2.4. Data Collection

Data were collected both on plot and plant bases from the three rows for all traits following the standard description of groundnut (IBPGR and ICRISAT, 1992). Plant basis data collection (pods per plant and seeds per pod), five plants were randomly taken and the mean values of these five plants were calculated.

#### 2.4.1. Phenological Data

**Days to 75% flowering (DTF) (days)**: It was recorded as the number of days from sowing to 75% of the plants in the plot started flowering.

**Days to 90% physiological maturity (DPM):** It was recorded as number of days from sowing to the stage when 90% of the plants in a plot have changed the color of their pods to dark.

**Grain filling period (GFP):** It was recorded as days from flowering to maturity, i.e. the number of days to maturity minus the number of days to 75% flowering.

#### 2.4.2. Yield and yield components

**Plant height (PH) (cm):** The mean height of main stem of five randomly taken plants was measured from the ground level or from the collar (point on the stem where roots start to grow).

**Number of branches per plant (NBP):** The mean number of branches per plant was obtained by counting the number of branches from each five sampled plants.

**Number of pods per plant (NPP):** This was determined as the mean value of five randomly sampled plants obtained by counting total number of pods per plant

**Number of seeds per pod (NSP):** The mean number of seeds per pod was obtained by counting the number of seeds collected from five mature pods from each five sampled plants.

**Pod yield (PY) (kg/ha):** The weight of dry pods from a unit plot was measured and was converted into kilogram per hectare.

**Shelling percentage (SP) (%):** This was calculated by dividing the weight of seeds to the weight of total pods shelled expressed as percent.

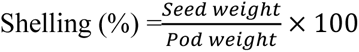

**Hundred-kernel weight (HKW**): Bulk of shelled seeds was counted and it was weighted and adjusted to standard moisture level (10%).

**Kernel yield (KY) (kg ha^-1^):** It was obtained from total harvest of the plot and adjusted to standard moisture level (10%) per plot in grams and converted into kilograms per hectare.

**Biomass yield (BY):** The weight in grams of sun dried above ground parts of the plants was recorded from the three rows and converted into kilogram per hectare.

**Harvest index (HI):** It was calculated as the ratio of kernel yield to total above ground biomass yield (biological yield).

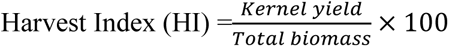

**Oil yield (OY):** It was calculated as oil content in percentage times kernel yield in ton/ ha.

### 2.5. Data Analysis

#### 2.5.1. Principal component analysis

Principal component analysis (PCA) was computed to find out the traits, which accounted more to the total variation. The data was standardized to mean zero and variance of one before computing principal component analysis to avoid differences in measurement scales. The principal component based on correlation matrix was calculated using SAS software version 9.0 (SAS, 2000).

#### 2.5.2. Genetic divergence and clustering of genotypes

Genetic distance of 36 groundnut genotypes was estimated using Euclidean distance (ED) calculated from quantitative traits after standardization (subtracting the mean value and dividing it by the standard deviation) as established by Sneath and Sokal (1973) as follows:

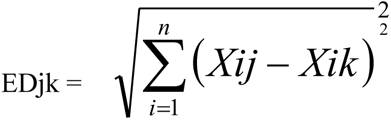

Where; EDjk = distance between genotypes j and k; xij and xik = phenotype traits values of the i^th^ trait for genotypes j and k, respectively; and n = number of phenotype traits used to calculate the distance. The distance matrix from phenotype traits was used to construct dendrogram based on the Unweighted Pair-group Method with Arithmetic Means (UPGMA). The results of cluster analysis were presented in the form of dendrogram.

## 3. Results

### 3.1. Principal Component Analysis

The principal component analysis (PCA) for 13 traits was computed to identify the critical traits which are important for the improvement of the crop and the traits that explained more of the variation in groundnut (Table 2). The results of PCA indicated that frst six principal components/factors were accounted for 74.50% of the total variance. Accordingly, the frst principal component (PC1) or factor 1 (F1) accounted for approximately 29.77% of the total variation which was influenced positively by quantitative characters viz. kernel yield (KY), oil yield (OY), dry pod yield (DPY), hundred seed weight (HSW), biomass yield (BY), number of branches per plant (NBP), plant height (PH), harvest index (HI), number of pods per plant (NPP) and shelling percentage (SP).

**Table 2.** Principal component values of the first six principal components from 13 traits of groundnut genotypes.

| Trait | PC1 | PC2 | PC3 | PC4 | PC5 | PC6 |
| --- | --- | --- | --- | --- | --- | --- |
| DTF | -0.07 | **0.90** | 0.07 | -0.05 | 0.13 | -0.03 |
| DTM | -0.08 | **0.94** | 0.003 | -0.05 | 0.13 | -0.001 |
| GFP | -0.10 | **0.56** | -0.24 | **0.40** | **-0.25** | **0.25** |
| NPP | **0.52** | -0.07 | 0.02 | **0.54** | 0.06 | **0.42** |
| PH | **0.55** | 0.17 | **0.40** | -0.08 | **0.36** | **0.32** |
| NBP | **0.66** | 0.04 | -0.04 | **0.41** | **-0.28** | -0.14 |
| BY | **0.68** | -0.005 | -0.07 | 0.20 | -0.12 | **0.26** |
| SP | **0.48** | -0.17 | **0.54** | **-0.23** | 0.06 | -0.02 |
| DPY | **0.79** | 0.16 | **-0.33** | -0.16 | 0.18 | -0.12 |
| KY | **0.91** | 0.05 | -0.22 | -0.21 | -0.10 | -0.13 |
| HSW | **0.76** | 0.24 | -0.09 | 0.06 | -0.10 | **-0.26** |
| HI | **0.54** | **-0.23** | **0.41** | 0.10 | **-0.37** | **-0.22** |
| OY | **0.89** | 0.03 | **-0.30** | **-0.24** | 0.08 | -0.10 |
| Eigenvalues | 5.04 | 2.50 | 1.73 | 1.33 | 1.04 | 1.00 |
| Proportions (%) | 29.77 | 14.71 | 10.16 | 7.82 | 6.14 | 5.90 |
| Cumulative (%) | 29.77 | 44.49 | 54.64 | 62.47 | 68.61 | 74.51 |

Principal component two (PC2) possessed an eigenvalue of 2.50 and showed 14.71% of total variation, which was largely influenced positively by only four quantitative characters viz. days to 75% flowering (DTF), days to physiological maturity (DTM), grain filling period (GFP) and hundred seed weight (HSW) whereas it was negatively influenced by HI in terms of factor loadings. This component discriminated the traits of genotypes based on phenology and productivity. An eigenvalue of 1.73 and 10.16% of variation was reflected by principal component three (PC3), which was influenced positively by quantitative characters viz. SP, HI and DPY. On the other hand, it was negatively influenced by OY in terms of factor loadings. Principal component four (PC4) contained eigenvalues of 1.33 and contributed 7.82% of the total variation. The traits which contributed more positively to PC4 were NPPS (0.54) followed by NBP (0.41) and (0.40) by GFP, while maximum negative value by OY (−0.24) followed by SP (-0.23). This component differentiated the genotypes based on phenology, productivity and quality. The fifth principal component (PC5) accounted for 6.14% of the total variation, with most of the variation being attributed to HI, PH and NBP. In PC6 highest positives were recorded for NPP (0.42), PH (0.32) and BY (0.26) on the other hand highest negative were recorded (−0.26) for HSW.

### 3.2. Genetic Divergence and Clustering of Genotypes

#### 3.2.1. Cluster analysis

According to the result of cluster analysis, groundnut genotypes (n=36) were grouped into six distinct clusters (Figure 1). Cluster-II comprised of 9 genotypes, which is about 25% of the total genotypes evaluated. Cluster III consisted eight (22.22%) of the total genotypes followed by cluster V which comprised seven (19.44%) genotypes. Cluster I consisted of six genotypes which is about 16.67% of the total genotypes tested. On the other hand, five genotype were included in clusters IV which accounted for 13.89 % whereas cluster V included only one genotype which is about 2.78% of total genotypes.

**Figure 1.**
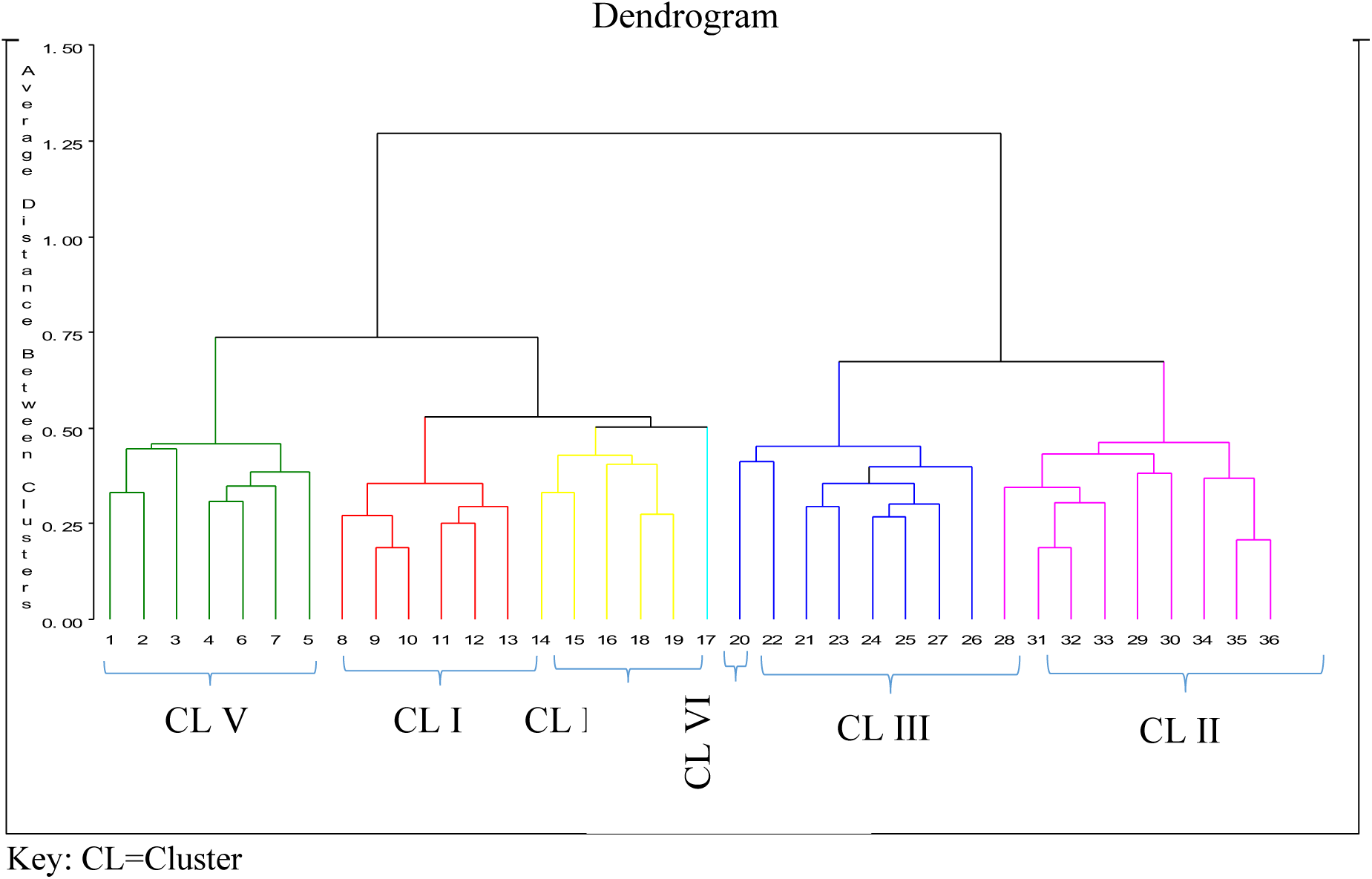
Dendrogram illustrating genetic divergence of groundnut genotypes on the basis of 13 traits.

#### 3.7.2. Cluster Mean Analysis

The mean values of four clusters for 13 quantitative traits are presented in Table 3. The result showed the differences between clusters by summarizing cluster means for the 13 traits. The highest cluster mean was recorded in cluster II for biomass yield per hectare (5620.53) and the lowest was recorded in cluster VI for OY (0.69). Most of the traits in cluster I had mean values lower than overall mean values, but had mean values greater than overall mean values of genotypes for HI.

**Table 3.** Mean values of six clusters for 13 traits of groundnut genotypes (n=36).

|  | CI | CII | CIII | CIV | CV | CVI | Overall mean |
| --- | --- | --- | --- | --- | --- | --- | --- |
| DTF | 55.33 | 55.22 | 54.88 | 57.40 | 57.64 | 55.0 | 55.93 |
| DTM | 114.58 | 115.39 | 114.94 | 117.80 | 118.21 | 116.0 | 116.06 |
| GFP | 58.58 | 57.44 | 60.06 | 60.60 | 58.86 | 62.50 | 59.07 |
| NPP | 36.18 | 37.77 | 39.80 | 31.62 | 33.41 | 50.80 | 36.62 |
| PH | 23.97 | 27.27 | 25.88 | 27.72 | 27.56 | 15.20 | 26.19 |
| NBP | 5.60 | 5.87 | 6.79 | 5.84 | 5.94 | 8.80 | 6.12 |
| BY | 4590.74 | 5620.53 | 5082.20 | 5420.00 | 4507.12 | 4469.4 | 5052.95 |
| SP | 65.67 | 69.22 | 64.63 | 69.30 | 66.43 | 61.00 | 66.85 |
| DPY | 3227.64 | 3622.89 | 3227.79 | 3659.62 | 3456.71 | 2254.8 | 3404 |
| KY | 2242.68 | 2616.96 | 2175.63 | 2631.85 | 2419.68 | 1446.7 | 2387.7 |
| HSW | 49.35 | 50.65 | 51.51 | 46.06 | 52.76 | 51.40 | 50.42 |
| HI | 47.23 | 45.82 | 43.17 | 49.77 | 53.52 | 30.83 | 47.09 |
| OY | 1.10 | 1.30 | 1.08 | 1.29 | 1.18 | 0.69 | 1.18 |

*DTF=days to flowering, DTM=days to maturity, GFP=grain filling period, NPP=number of pods per plant, PH=plant height, NBP=number of branches per plant, BY= biomass yield, SP=shelling percentage, DPY=dry pod yield, KY=kernel yield, HSW=hundred seed weight, HI=harvest index, OY=oil yield.*

The groundnut genotypes in cluster II showed highest mean value for most of the studied traits compared with their respective overall mean. It had mean values higher than overall mean values for all traits, but had mean values lower than overall mean values of genotypes for DTF, DTM, GFP, NBP and HI. Cluster III had mean values greater than overall mean values of genotypes for GFP, NPP, NBP, BY, and HSW, while for DTF, DTM, PH, SP, DPY, KY and OY it had mean values lower than overall mean values.

On the other hand, cluster IV contained greater mean values than overall mean values for DTF, DTM, GFP, PH, BY, SP, DPY, KY, HI and OY, but had lower mean values than overall mean values for NPP, NBP and HSW. Traits such DTF, DTM, PH, DPY, KY and HI are contained greater mean values than overall mean value in cluster V. Majority of traits in the cluster VI are contained lower mean values than overall mean. GFP, NPP, NBP and HSW had greater mean values than overall mean of individual traits.

#### 3.7.3. Genetic Distance among Groundnut Genotypes

The genetic distances for all possible pairs of 36 groundnut genotypes are presented in Appendix Table 1. The genetic distances of genotypes varied from 2.45 to 8.54, with 5.44, 1.17, and 21.56% for the mean, standard deviation, and coefficient of variation, respectively (Table 4). The highest Euclidean distance between the groundnut genotypes was observed between Gv17 and Gv28 (8.54), followed by Gv3 and Gv23 (8.49), Gv3 and Gv30 (8.49), Gv15 and Gv17 (8.37), Gv22 and Gv28 (8.37) and Gv3 and Gv34 (8.36) (Appendix Table 1). While, the lowest genetic distance was found between Gv13 and Gv24 (2.45), and Gv14 and Gv36 (2.45), followed by Gv18 and Gv31 (2.65), Gv9 and Gv10 (2.83), Gv10 and Gv27(8.83), and Gv31 and Gv32 (2.83) (Appendix Table 1).

**Table 4.**
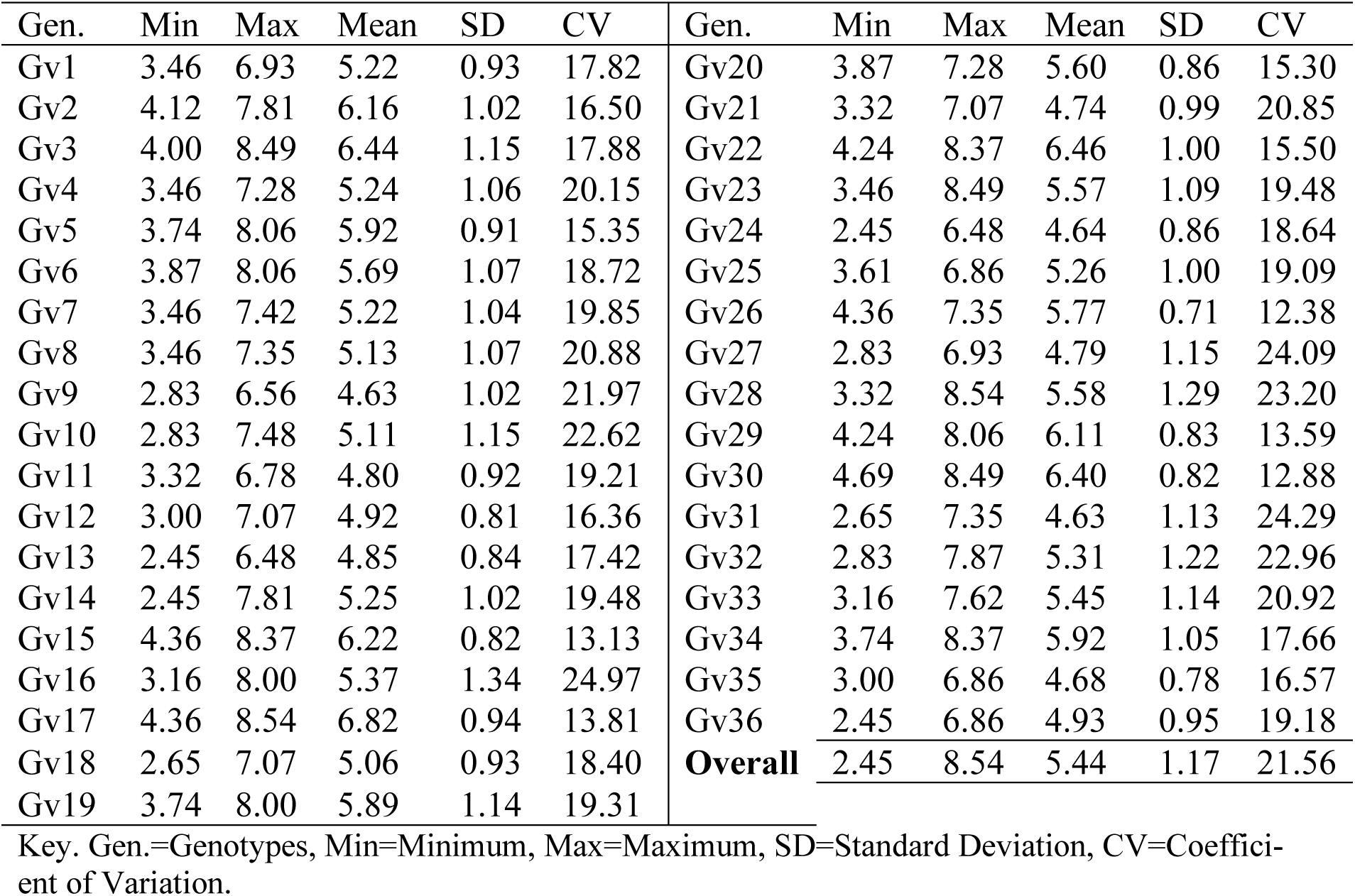
Range and mean Euclidean distance of groundnut genotypes estimated from 13 quantitative traits (n=36).

The maximum Euclidean distance was obtained for genotype Gv17 (6.82) followed by Gv22 (6.46), Gv3 (6.44) and Gv30 (6.40), which also had Euclidean distance greater than the overall mean (5.44). Totally, 17 genotypes exhibited Euclidean distance greater than overall mean.

## 4. Discussion

Multivariate analyses, viz. principal component analysis, UPGMA clustering and genetic distance analysis, were performed on a set of 36 groundnut genotypes by 13 traits, viz. days to flowering, days to physiological maturity, grain filling period, number of branches per plant, number of pods per plant, plant height, biomass yield, shelling percentage, dry pod yield, hundred seed weight, harvest index, kernel yield and oil yield.

Grouping genotypes based on their agro-morphological and quality characters is useful as it assists in identification and selection of best performers and genetically diverse parents for use in breeding program (Govindaraj, 2015; Niveditha *et al.,* 2016). In the current study the principal component analysis revealed six components with eigenvalues greater than a unit. Principal components with eigenvalues greater than one are theoretically have more information than any single variable alone (Iezzoni & Pritts, 1991). The first and the second component contributed most of the variation among the total variations explained. Similar results were reported in groundnut (Mubai *et al.,* 2020; Seltene *et al*. 2021). The first component had eigenvalue of 5.04 and was effective in partitioning yield, yield related and quality traits. This component can be called productivity and quality since they discriminated the genotypes according to their yield and quality. The second component was associated with DTF, DTM, GFP and discriminated the genotypes on the basis of maturity duration. This is due to the fact that effective discrimination of genotypes into different components through relatively higher contribution of few characters rather than small contribution from each character (Chahal and Gosal, 2002 and Yan and Tincker, 2005).

The study indicated the presence of diversity among the tested groundnut genotypes. Groundnut genotypes grouped in different clusters could be evaluated for combining ability to constitute a pool of best parents. These findings are supported by previous reports by Mohammed *et al*. (2022); and Khan *et al.,* 2020 that there is high genetic diversity in groundnut. The grouping of the genotypes indicated that evaluated characters had influence on clustering pattern. Similar result was reported in groundnut by Seltene *et al*. (2021); and Makinde and Ariyo. (2010).

The genetic distance of for all possible pairs of 36 groundnut genotypes ranged from 2.45 to 8.54 with the mean, standard deviation and coefficient of variation of 5.44, 1.17 and 21.56%, respectively. A total of 17 (47.2%) out of 36 genotypes had mean genetic distances greater than the overall mean genetic distance of genotypes. These results were supported by previous findings of Bonny *et al*. (2019) and Mubai *et al*. (2020) who reported six clusters and variations of genetic distance among one hundred and one Bambara groundnut germplasms. These results suggested that the genotypes that had greater mean of genetic distance over the average mean distance were more distant to others and crossing between these genotypes could be used to combine desired traits in progeny of subsequent generation. Thus, the maximum amount of heterosis, which could have implication for genetic improvement of the crop for the target trait, is expected from the crosses with genotypes that are distant from each other. Based on the result of the genetic distance analysis using Euclidean method, there were variations of genetic distances among genotypes, Gv17 and Gv28, Gv3 and Gv23, Gv3 and Gv30, Gv15 and Gv17, Gv22 and Gv28, and Gv3 and Gv34 which could be exploited through hybridization for cultivar development in groundnut breeding programs in Ethiopia. Therefore, these genotypes are recommended as excellent candidates for further breeding and variety development.

## Declaration of Computing Interest

The authors declare that none of the work reported in this study could have been influenced by any known competing financial interests or personal relationships.

## Data availability

The data availability was requested through the author’s email.

## Author statement

Statement of contribution of Desu Beriso was done original study, design methodology, data retrieval, software, data analysis and interpretation, writing original draft. Dr Seltene Abady and Dr Abdi Mohammed were writing curative data analysis, interpretation and editing the review article.

## Acknowledgment

We are highly acknowledged Dire Dawa and Tony farm Research sub-Station of Haramaya University for unreserved support for provided me field experiment to conducting research article. The authors also expressed gratitude to two anonymous reviewers for their thoughtful comments.

## Appendix

**Appendix Table 1.**
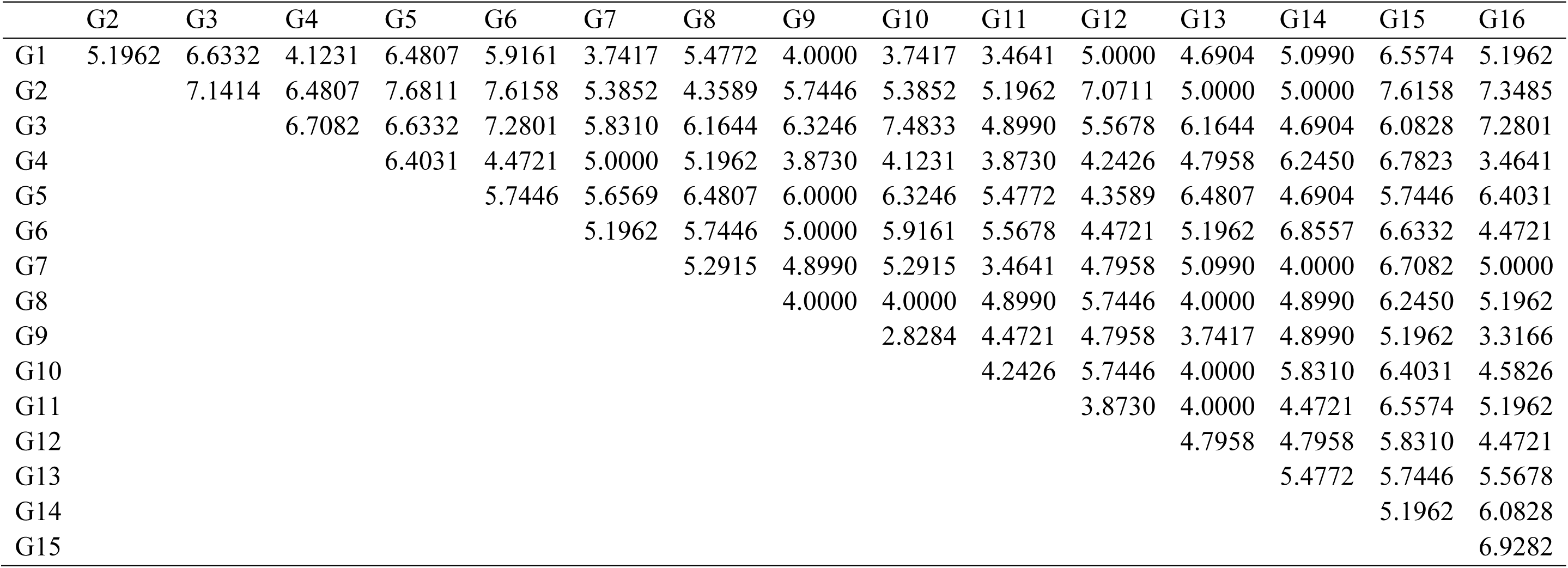
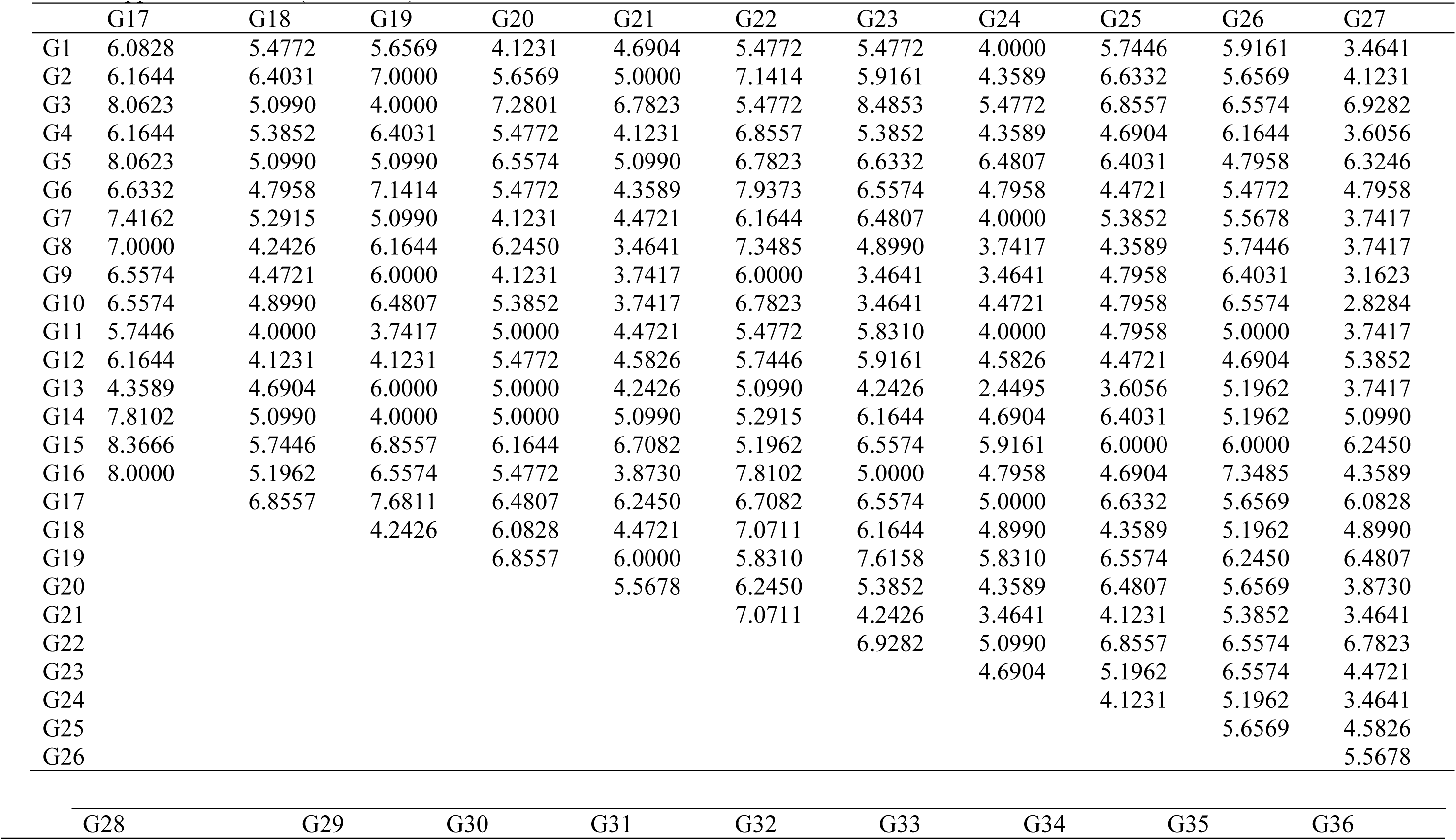
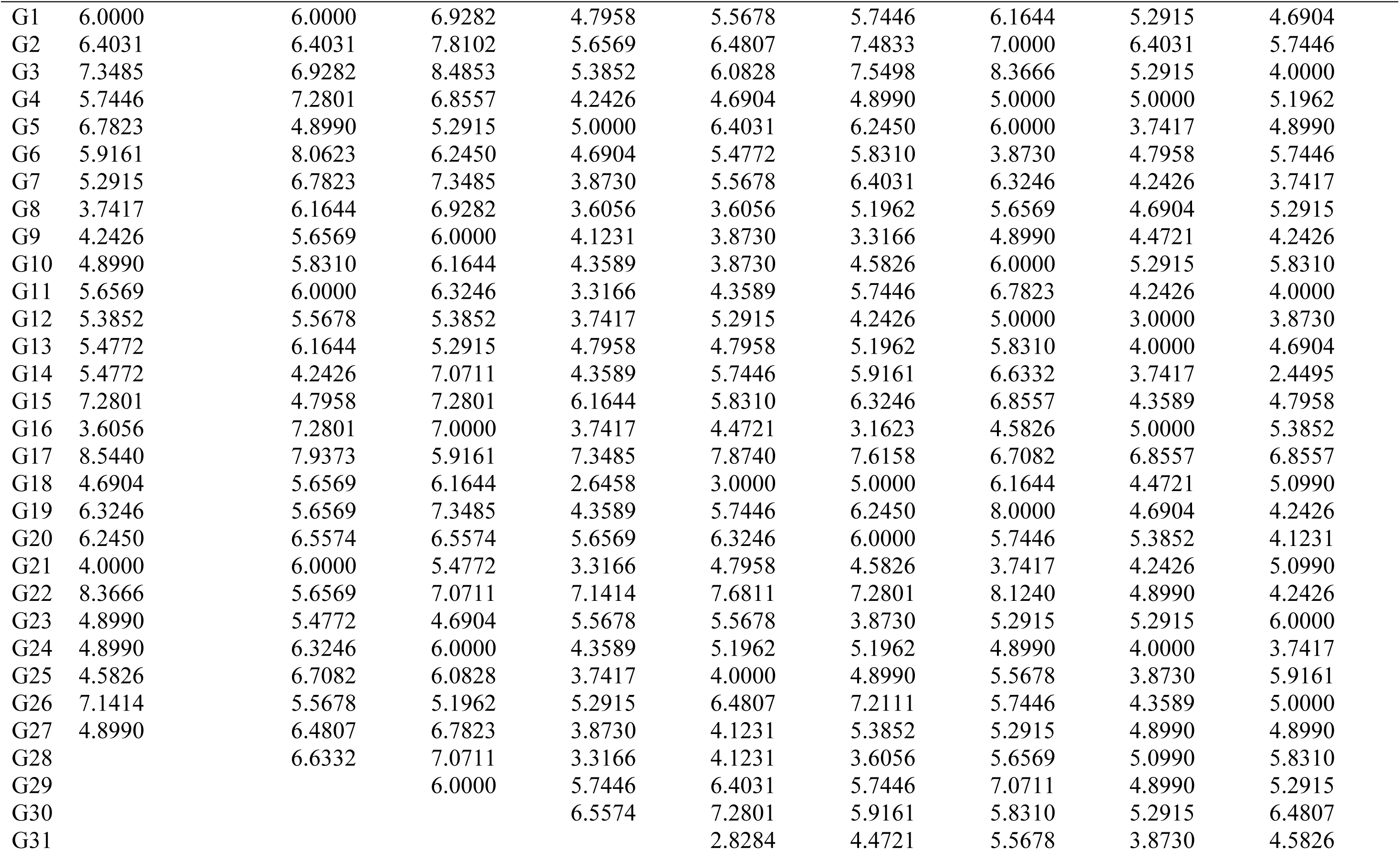
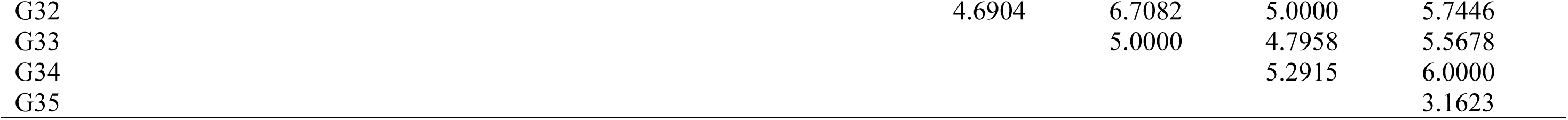
Euclidean distances of groundnut genotypes estimated from mean values of genotypes for 13 quantitative traits (n=36).

